# Effect of oxidized β-carotene-oxygen copolymer compound on health and performance of pre- and post-weaned pigs

**DOI:** 10.1101/2020.08.07.241174

**Authors:** La Van Kinh, William W. Riley, James G. Nickerson, Doan Vinh, Nguyen Van Phu, Nguyen Thanh Van, La Thi Thanh Huyen, Graham W. Burton

## Abstract

The discovery that a naturally occurring, biologically active β-carotene-oxygen copolymer compound is the main product formed in spontaneously oxidized β-carotene has stimulated interest in its potential health benefits. The copolymer, formed in nature or synthetically by the air-oxidation of β-carotene, possesses beneficial immune modulating activities that previously had been attributed to β-carotene itself. Support for these benefits is provided by previous studies showing that supplementation in feed with low parts-per-million levels of copolymer-rich, fully oxidized β-carotene (OxBC) helped reduce the negative impact of subclinical necrotic enteritis in broilers and improved growth in weaned piglets. To further assess these potential benefits, two trials were conducted in swine raised under commercial conditions in Vietnam. Trial 1, a 140-day full-grow, post-wean study with 500 28-day-old pigs, compared 2, 4 or 8 ppm OxBC against both an unsupplemented and an antibiotic control group. OxBC and antibiotics each improved growth rate, feed efficiency, and body weight compared to the control (P<0.001). Animals receiving 4 and 8 ppm OxBC performed better than did animals on antibiotics (P<0.001). In starter pigs, OxBC reduced the occurrence of diarrhea dose-dependently (4 and 8 ppm) and to a greater extent than did antibiotics (P<0.001). Trial 2, a 49-day study with 420 piglets, was conducted in two-stages. In Stage 1 (pre-wean), OxBC in the transition (creep) feed produced a dose-dependent trend toward increased body weight over 21 days, reaching significance at the highest inclusion level (16 ppm) (P<0.001). In Stage 2 (post-wean), body weight gain showed a dose-dependent trend and was significant for both 8 ppm OxBC and the antibiotics at 28 days post-wean (P<0.001). Feed conversion was better at 8 ppm OxBC and for the antibiotic group (P<0.001). These findings support the concept that β-carotene-oxygen copolymers help optimize immune function, and provide validation for the effectiveness of this strategy in enhancing animal performance in the absence of in-feed antibiotics.

## Introduction

We have previously reported that the spontaneous non-enzymatic oxidation of β-carotene produces predominantly an oxygen-rich copolymer compound with immunomodulatory properties (Burton et al., 2014; Johnston et al., 2014). Fully oxidized β-carotene, termed OxBC and rich in copolymer compound, exerts its actions on the immune system through pathways that are distinct from either vitamin A or intact β-carotene, which are both absent. We have proposed that copolymer compounds are in fact the actual agents responsible for many of the provitamin A-independent activities of β-carotene and other carotenoids (Johnston et al., 2014).

OxBC exhibits dual immunological activities relating to: a) enhancing innate immune detection and response to pathogens (Johnston et al., 2014) and, b) an anti-inflammatory/pro-resolution action that limits the extent of over-zealous immune responses and reduces the level of background inflammation (Duquette et al., 2014; Chen et al., 2020).

The utility of OxBC as a feed additive and alternative to antibiotic growth promoters has been demonstrated in studies with piglets (Hurnik et al., 2011), sows (Chen et al., 2020) and broiler chickens (Kang et al., 2018). In piglets, dietary supplementation with OxBC improved growth performance and prevented the vaccine-induced growth-lag associated with the PRRS (porcine reproductive and respiratory syndrome) vaccination (Hurnik et al., 2011). In sows, supplementation with OxBC, begining at late gestation and continuing through lactation, resulted in reduced proinflammatory cytokine levels in colostrum and milk concurrent with increased colostral and milk immunoglobulin levels (Chen et al., 2020). In broilers, dietary supplementation with OxBC reduced the level of pathogen (*Clostridium perfringens*) recovered from the gut and protected against the reduction in growth performance associated with induction of subclinical necrotic enteritis (Kang et al., 2018).

Several authors have proposed that the search for suitable alternatives to antibiotic growth promoters should focus on substances that achieve effects similar to the antibiotics they are intended to replace, namely, reduction in both bacterial load and inflammation (Niewold, 2007, 2010; Khadem et al., 2014; Soler et al., 2016). The results from trials with pigs and poultry highlight the utility of OxBC in achieving both of these outcomes. The benefits observed with piglets and gestating/lactating sows likely are consistent with the anti-inflammatory actions of OxBC, whereas, the reduction in *C. perfringens* in the poultry study supports the benefits of innate immune priming. Note that OxBC has no direct anti-microbial effect and that the reduction in *C. perfringens* in the poultry study reflects actions on the host’s immune system, which is better able to detect and respond to the presence of pathogens.

The study objective was to determine the effects of OxBC on the performance of swine through their full growth period under commercial production conditions in Vietnam. The first trial evaluated OxBC over an entire 140-day post-wean growth cycle., while the second trial evaluated the effect of OxBC on the health and growth performance of pre-wean and post-wean pigs. We hypothesized that OxBC would serve as a an effective substitute for in-feed antibiotics in real-world, commercial, swine-production conditions.

## Materials and Methods

All animal care procedures followed the procedure approved by the Animal Care and Use Committee of the Institute of Animal Science for Southern Vietnam. These procedures approved by the committee were established in accordance with Vietnam’s Law on Animal Husbandry, which covers the humane treatment of livestock in transport, slaughter, scientific research and other activities.

### Trial 1. Full grow study

The trial was carried out at a commercial farm, Thai My Pig farm, Thai My commune, in the Cu Chi district, Ho Chi Minh City, from 17 May to 30 November 2014.

Five hundred weaned barrows and gilts (28 days of age) were used in a 140-day complete wean-to-finish feeding trial. The animals were reared on site and were the progeny of Landrace x Yorkshire females and Duroc boars. The herd was asymptomatic for PRRS and was vaccinated against hog cholera (Coglapest™, Ceva-Phylaxia Veterinary Biologicals Co. Ltd, Hungary), foot-and-mouth disease (Biotaftogen™, Biogénesis Bagó, Argentina) and PRRS (Tai Xanh™, Navetco, Vietnam).

Animals were randomized by weight with each pen containing 20 pigs with the same ratio of barrows to gilts. The trial was comprised of five dietary treatment groups with five replicate pens per treatment as follows: Control (basal diet with no antibiotics or OxBC), AB (basal diet with antibiotics; no OxBC) and OxBC (basal diet supplemented with 2, 4 or 8 ppm (mg/kg) OxBC; no antibiotics). The compositions of the basal diets for the Starter, Grower and Finisher phases of the study are shown in Table 1. No in-feed or water-administered medications or feed additives were employed with the exception of the antibiotics used in the AB group and OxBC used in the three OxBC groups. OxBC was provided in-feed as the OxC-beta™ Livestock 10% commercial premix product (Avivagen Inc. Ottawa, Canada). Chlortetracycline and colistin sulfate were purchased from Zhumadian Huazhong Chia Tai Co. Ltd (China) and Lifecome Biochemistry Co. Ltd (China), respectively.

**Table 1.**
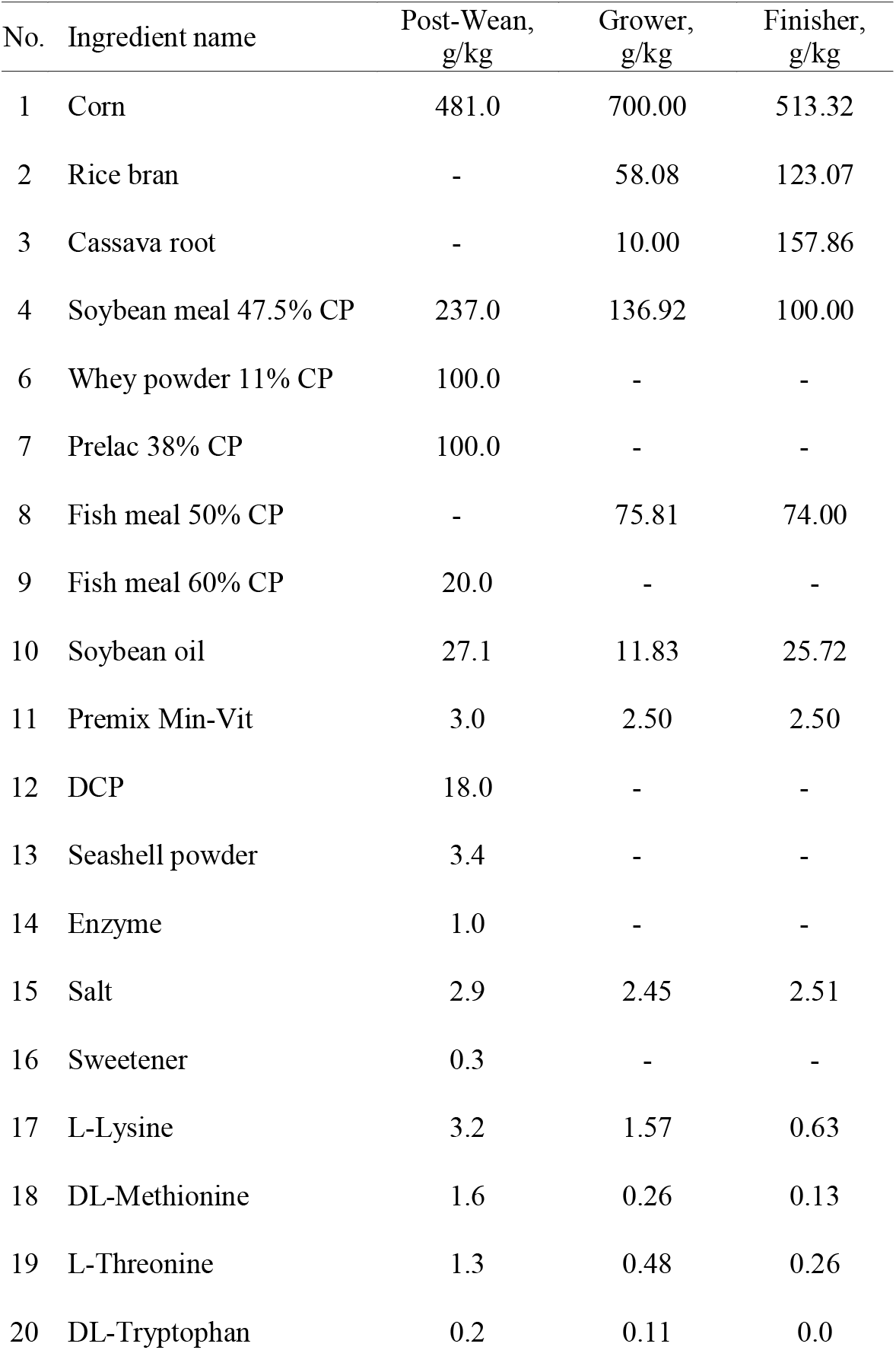
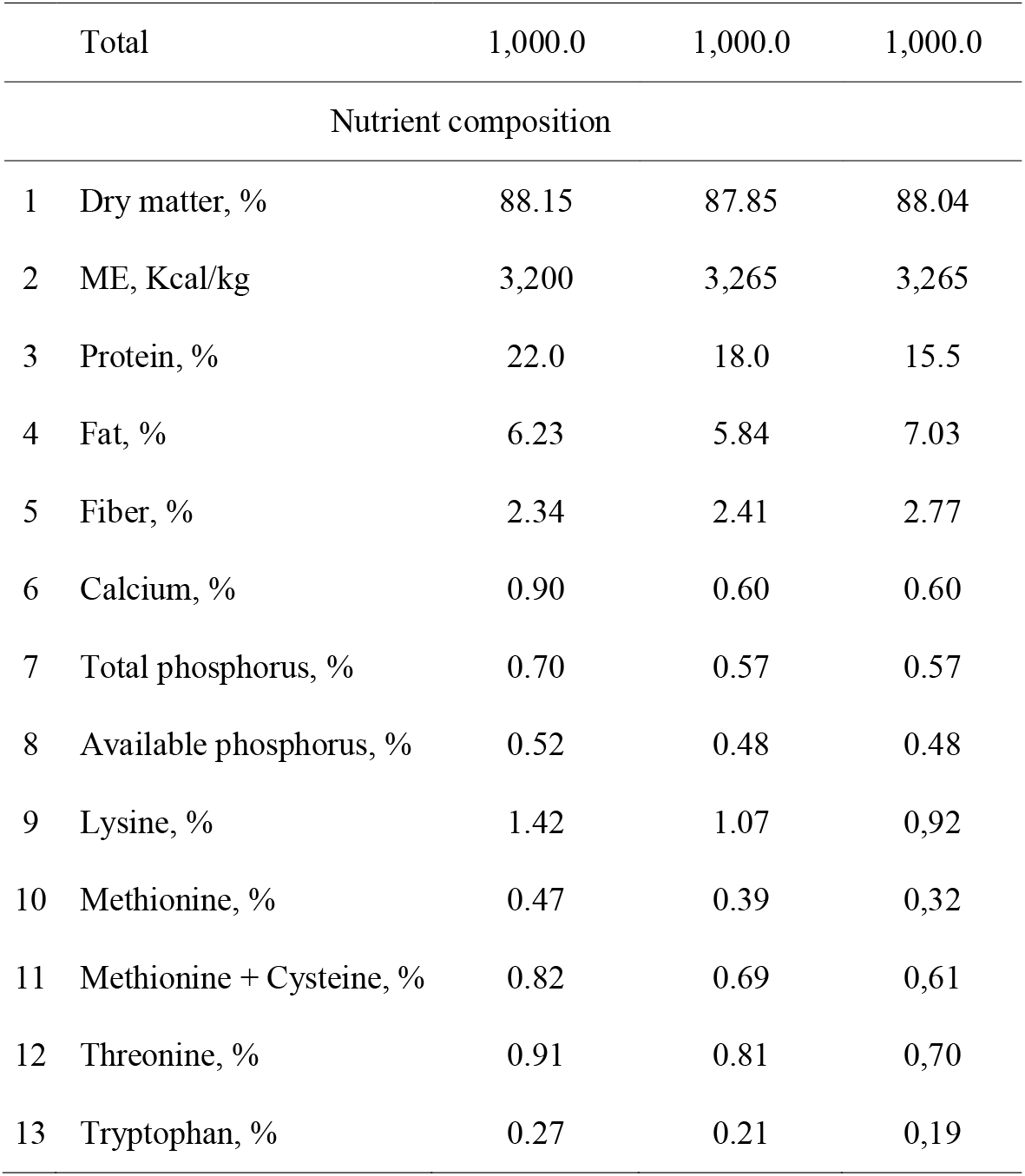
Composition of basal diets for Starter, Grower and Finisher phases of the full grow study

For logistical and space purposes, the trial was conducted in a time-replicated fashion with replicates one and two for all five treatment groups beginning on Day 1; replicates three and four for all five treatment groups began the trial 28 days later, and replicate five for all five treatment groups began the trial 56 days after the first cohort.

Each pen was equipped with a dry hopper feeder with six feed access holes, each measuring 13 cm x 13 cm, aligned parallel to the pen front. Feeders were located at the front of the pen, and a water drinker was located at the back of each pen. The pigs had *ad libitum* access to feed and water throughout the trial.

Pigs were individually weighed at the start of the study (Day 1) and at Days 28 (~20 kg), 84 (~50 kg) and 140 (~100 kg). All feed given was weighed daily, the feeders were emptied weekly and the remaining feed weighed.

Growth performance parameters were calculated for each stage of the production cycle (Starter: Days 1-28, Grower: Days 29-84, and Finisher: Days 85-140) as well as for the overall study period (Days 1-140).

Statistical analyses were performed using the Statistical Analysis System for Personal Computers (SAS) Version 9.1 (SAS Institute, Cary, NC). Treatment effects were tested with a combination of analysis of variance for continuous variables using the MIXED procedure and Chi-square analysis, including the Cochrane-Armitage trend test. Dichotomous variables (pigs clinically treated or removed) were created using the post-mortem and clinical results from the data provided to analyze the number of pigs removed from or treated during the study. The data were analyzed by logistic regression using the GLIMMIX procedure.

### Trial 2. Pre-and post-wean piglet study

The trial was carried out at a commercial farm, Thai My Pig farm, Thai My commune, in the Cu Chi district, Ho Chi Minh City from March 15 to May 30, 2016.

Forty-two (42) sows and their offspring were used in the two-stage, 49-day feeding trial. Stage 1 evaluated the benefits of supplementation with OxBC in creep feed in pre-weaned piglets. Stage 2 evaluated the benefits of continuing OxBC supplementation through the post-wean Starter phase.

All piglets were the progeny of Landrace x Yorkshire females and Duroc boars and were reared on site. The piglets were vaccinated against PRRS (Tai Xanh™, Navetco Vietnam) at 14 days old.

After one day post-farrowing, sows with minimum litter sizes of ten piglets and similar weights were selected. Litter sizes were also standardized by cross-fostering between sows within treatment groups within three days post-farrowing. As most sows had litter sizes greater than ten, surplus piglets were cross-fostered to sows that were not part of the trial.

#### Stage 1. Pre-wean (Days 0-21)

In Stage 1, 42 lactating sows (one week post-farrow) and their one-week-old offspring were randomly and evenly assigned to one of six treatments with seven replicate pens per treatment. Each pen contained one sow and her litter of ten piglets. The mean piglet body weight was 2.44 kg. As there were insufficient sows to carry out seven replicates simultaneously, the trial was run in a time-replicated fashion, with two cohorts of sows. The first cohort contained 18 sows with 180 piglets (six treatments and three replicates) and the second cohort contained 24 sows with 240 piglets (six treatments and four replicates). The second cohort began the trial one week after the first cohort.

The diets were introduced as soon as the piglets were ready to begin consuming creep feed. Sows in all groups received a basal commercial lactation diet without OxBC supplementation. Dietary treatments were delivered to piglets via supplementation of the creep feed as follows: Control (basal diet with no antibiotic or OxBC), AB (basal diet with antibiotics) and OxBC (basal diet with 2, 4, 8 or 16 ppm OxBC; no antibiotic). The composition of the basal diet is shown in Table 2.

**Table 2.** Composition of the basal diets for the Creep and Starter phases of the pre- and post-wean study.

| Ingredient Name | Pre-Wean, g/kg | Post-Wean, g/kg |
| --- | --- | --- |
| Corn | 573.2 | 636.5 |
| DDGS | - | 30.0 |
| Soybean meal 47% CP | 120.0 | 200.0 |
| Soya oil | 30.0 | 31.5 |
| Skimmed milk replacer | 93.7 | - |
| Whey powder | 50.0 | - |
| Blood plasma protein | 43.0 | 39.3 |
| Fish meal | 70.0 | 33.5 |
| Dicalcium phosphate | 5.6 | 09.9 |
| Calcium carbonate | 02.5 | 06.8 |
| Salt | - | 02.5 |
| Vitamin-mineral premix | 5.0 | 03.0 |
| L-Lysine HCl | 2.7 | 03.6 |
| Threanine | 1.0 | 1.0 |
| Methionine | 1.8 | 1.6 |
| Tryptophan | 1.2 | 0.8 |
| Sweetener | 0.3 | - |
| Total | 1000 | 1000 |
| Chemical composition |  |  |
| Crude protein, % | 22.21 | 21.29 |
| Crude fiber, % | 2.01 | 2.85 |
| Crude fat, % | 6.32 | 6.42 |
| NE swine, kcal/kg | 2557 | 2486 |
| ME swine, kcal/kg | 3275 | 3271 |
| Calcium, % | 0.85 | 0.75 |
| Phosphorus, % | 0.70 | 0.65 |
| Available phosphorus, % | 0.47 | 0.40 |
| Sodium, % | 0.29 | 0.28 |
| Lysine, % | 1.42 | 1.32 |
| Methionine, % | 0.54 | 0.45 |
| Cysteine, % | 0.30 | 0.32 |
| Methionine+Cysteine, % | 0.85 | 0.79 |
| Threonine, % | 0.89 | 0.83 |
| Tryptophan, % | 0.31 | 0.29 |
| Arginine, % | 1.10 | 1.18 |
| Isoleucine, % | 0.79 | 0.73 |
| Leucine, % | 1.74 | 1.68 |
| Valine, % | 0.97 | 0.90 |
| Histidine, % | 0.52 | 0.52 |
| Phenylalanine, % | 0.92 | 0.92 |

Feed and water were provided *ad libitum*. Piglets were individually weighed at the start of the study (Day 1) and at weaning (Day 21). Feed given was weighed daily and unused feed was weighed at the end of each week.

#### Stage 2. Post-wean (Days 22-49)

Stage 2 began at Study Day 22, one day post-wean, when the piglets were 29 days old. Three hundred and sixty (360) piglets were used in a random block design of six treatments with six replicate pens of ten piglets each. Piglets were randomized within their prior treatment groups before being equally assigned into pens. Treatments were as follows: Control (basal diet with no antibiotics or OxBC), AB (basal diet with antibiotics, no OxBC) and OxBC (basal diet with 2, 4, 8 or 4 (16 in Stage 1) ppm OxBC; no antibiotics). The composition of the basal diet is shown in Table 2. The piglets were provided feed and water *ad libitum*. Feed given was weighed daily and any unused feed was weighed at the end of each week.

The following measurements were performed: mean body weight (BW) at Days 1, 21 and 49 for each animal; average daily feed intake (ADFI): total feed intake per pen divided by the product of the number of live animals and days on diet; ADG: total BW gained per pen divided by the product of the number of live animals and days on diet; FCR: total feed consumption divided by BW increase, defined as: (sum of final body weights of surviving animals plus weight of mortalities and removals) – (sum of initial body weights including initial bodyweight of animals dead or removed during the specified period); mortality and diarrhea incidence, for each treatment group for both Stage 1 and Stage 2 of the study. Diarrhea rate was determined as the percentage of animals presenting with loose or watery stools.

Data were analyzed by ANOVA using Minitab Statistical Software (Minitab, State College, PA, USA). Differences between means were tested using Fisher’s multiple range test when the F value was significant at P< 0.05. Data were considered significantly different at P<0.05.

## Results

### Trial 1. Full grow study

The effect of dietary supplementation with OxBC on the growth performance of pigs is shown in Table 3. On Day 1, there were no significant differences in BWs among any of the groups (P>0.05). Dietary supplementation with all three levels of OxBC during the Starter period (Days 1 to 28) resulted in significant improvements in body weight (BW), average daily weight gain (ADG), average daily feed intake (ADFI) and feed/gain ratio (F/G) compared to the control group (P<0.001). Animals in the OxBC groups also performed better than or equivalent to those receiving antibiotics during the starter period.

**Table 3.**
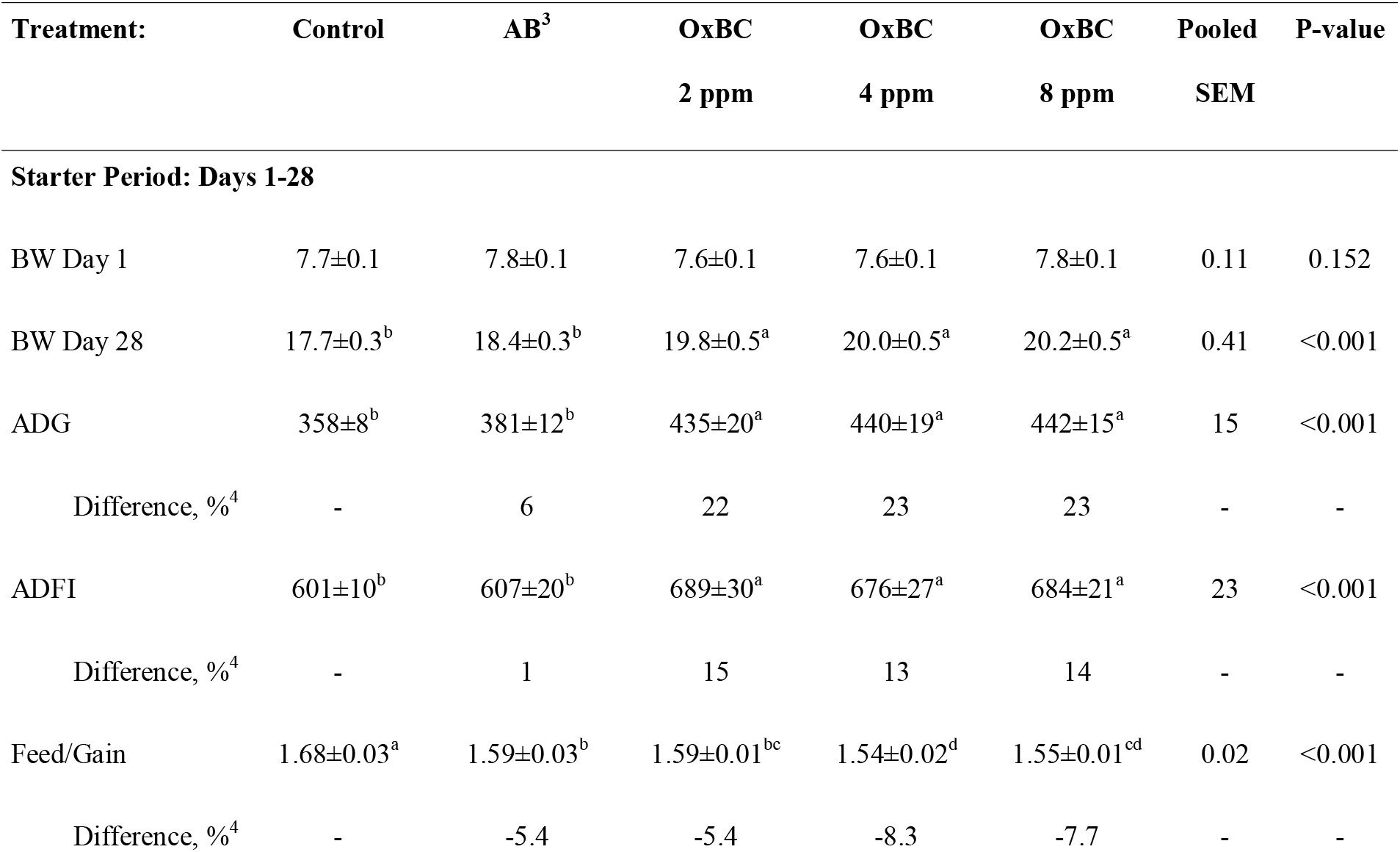
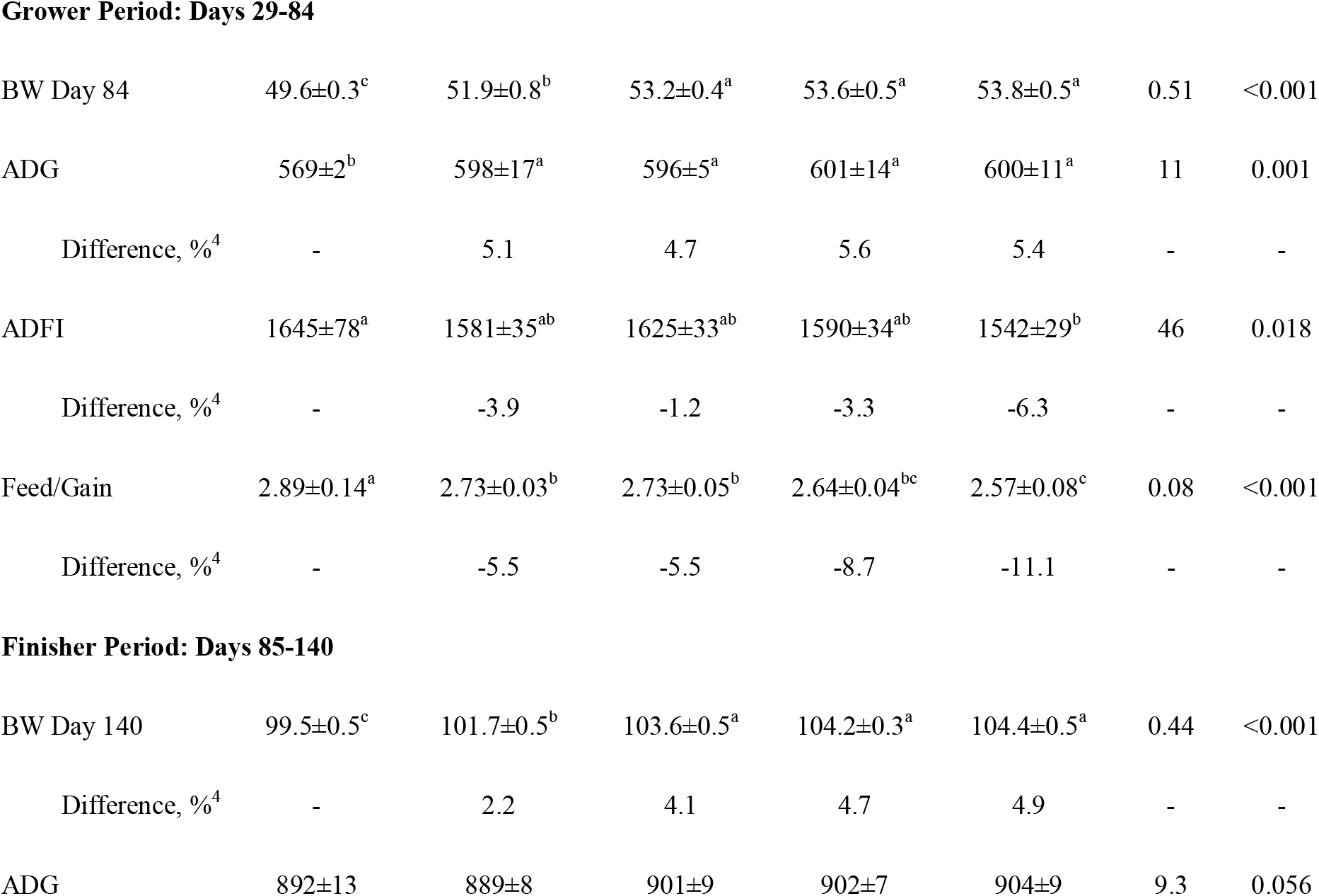
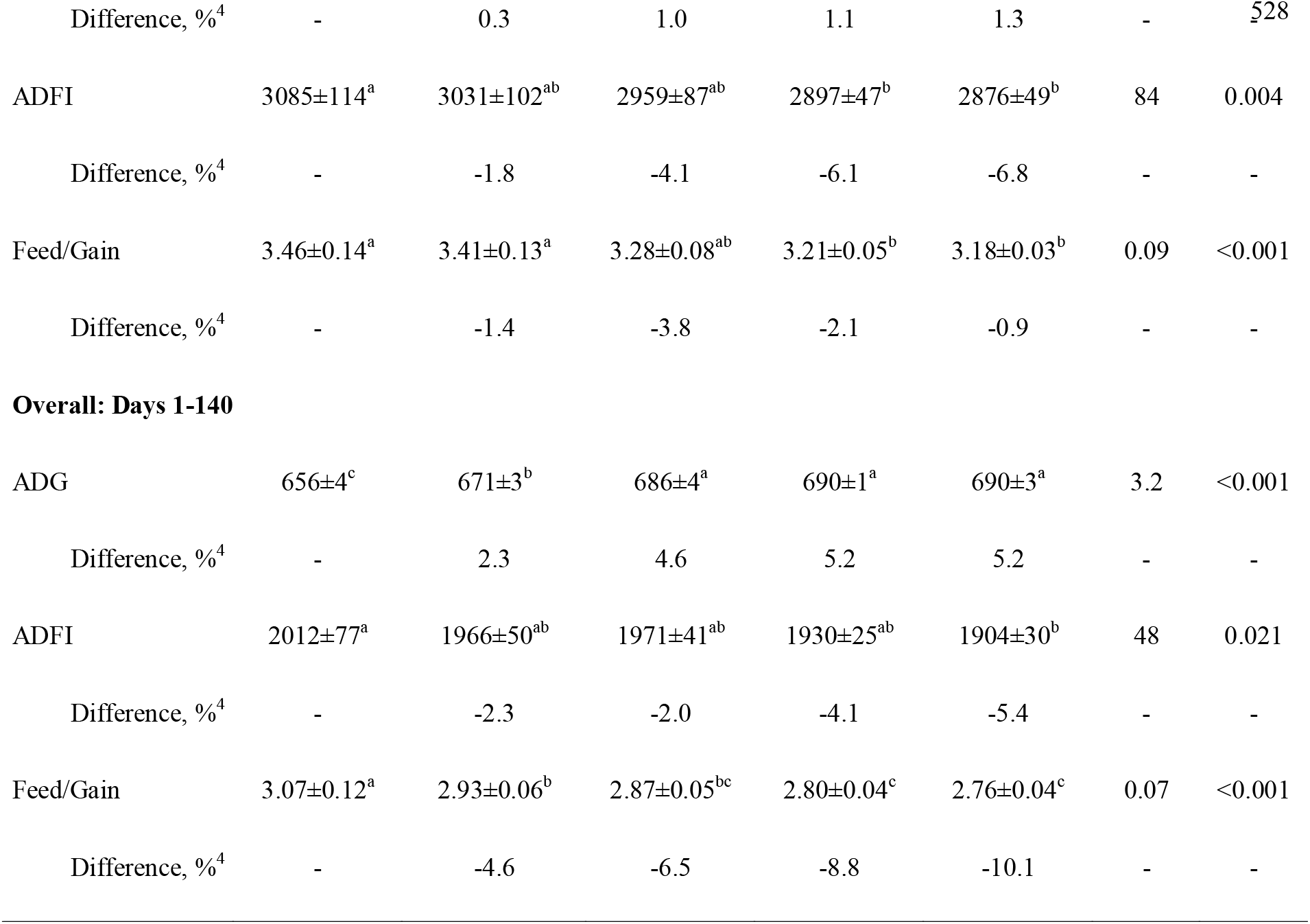
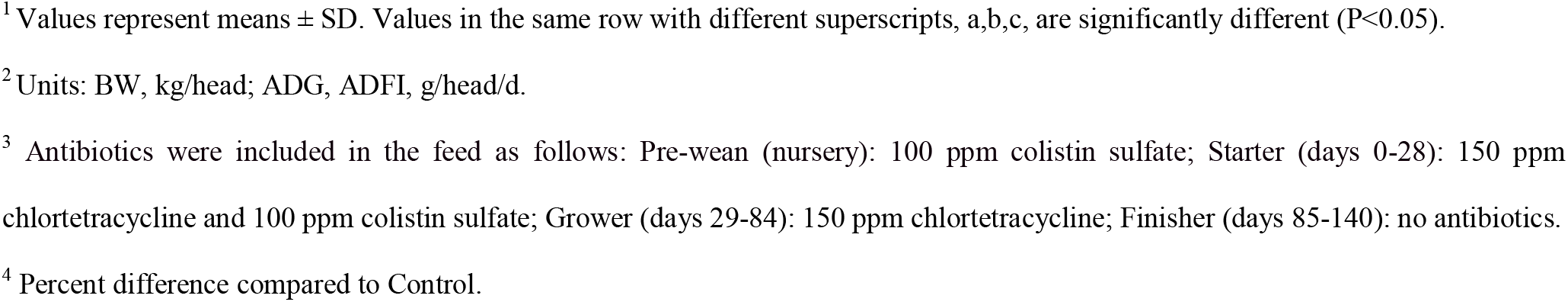
Effect of dietary OxBC on ADG, ADFI, F/G and BW of pigs in the Starter, Grower, Finisher and Overall periods.^1,2^

During the Grower period (Days 29-84), supplementation with all levels of OxBC significantly increased ADG relative to the Control (P<0.001). The ADG of the AB group was also higher than the Control (P<0.001), but there were no differences among the OxBC and the AB groups. There was an apparent dose-dependent decrease in feed consumption across the three OxBC groups, with the ADFI of the 8 ppm OxBC group being significantly lower than the Control (P<0.05). The F/G ratios for all OxBC groups and the AB group were significantly improved compared to the Control (P<0.001). The 8 ppm OxBC group had the lowest F/G ratio, which was significantly lower than the ratios for the Control, AB and 2 ppm OxBC groups (P<0.001).

During the Finisher stage (Days 85-140), the ADGs of animals in the OxBC groups were greater than those of either the Control or AB groups, although the differences did not reach statistical significance (P>0.05). Supplementation with 4 or 8 ppm OxBC resulted in significant improvements in the F/G ratio compared to the Control and AB groups (P<0.001). As was observed in the grower period, animals receiving OxBC had lower feed intake during the finisher phase when compared to the Control. The feed intake was significantly lower (P<0.005) in the 4 ppm OxBC and 8 ppm OxBC groups compared to the Control group. Note that during the finisher stage, animals in the AB group did not receive antibiotics, which is a common practice and legal requirement in many jurisdictions, including Vietnam.

Overall (Study Days 1-140), dietary supplementation with OxBC at 2, 4 or 8 ppm led to significant increases in final BW and ADG, relative to both the Control and the AB groups (P<0.001). OxBC also reduced the overall F/G (P<0.001) and ADFI (P<0.05) compared to the Control with the 4 and 8 ppm groups also outperforming the AB group on F/G (P<0.001).

In terms of clinical health, the results presented in Table 4 show that the highest incidences of both diarrhea and mortality were observed in the Control group for all study periods, with the lowest being observed in the OxBC groups. The highest incidence of diarrhea was during the Starter period, and OxBC significantly and dose-dependently reduced the diarrhea rate (P<0.001). Diarrhea was the lowest (less than half that of the Control group) in the 8 ppm OxBC group, and both the 4 and 8 ppm OxBC groups had significantly reduced incidences relative to the AB group (P<0.001).

**Table 4.**
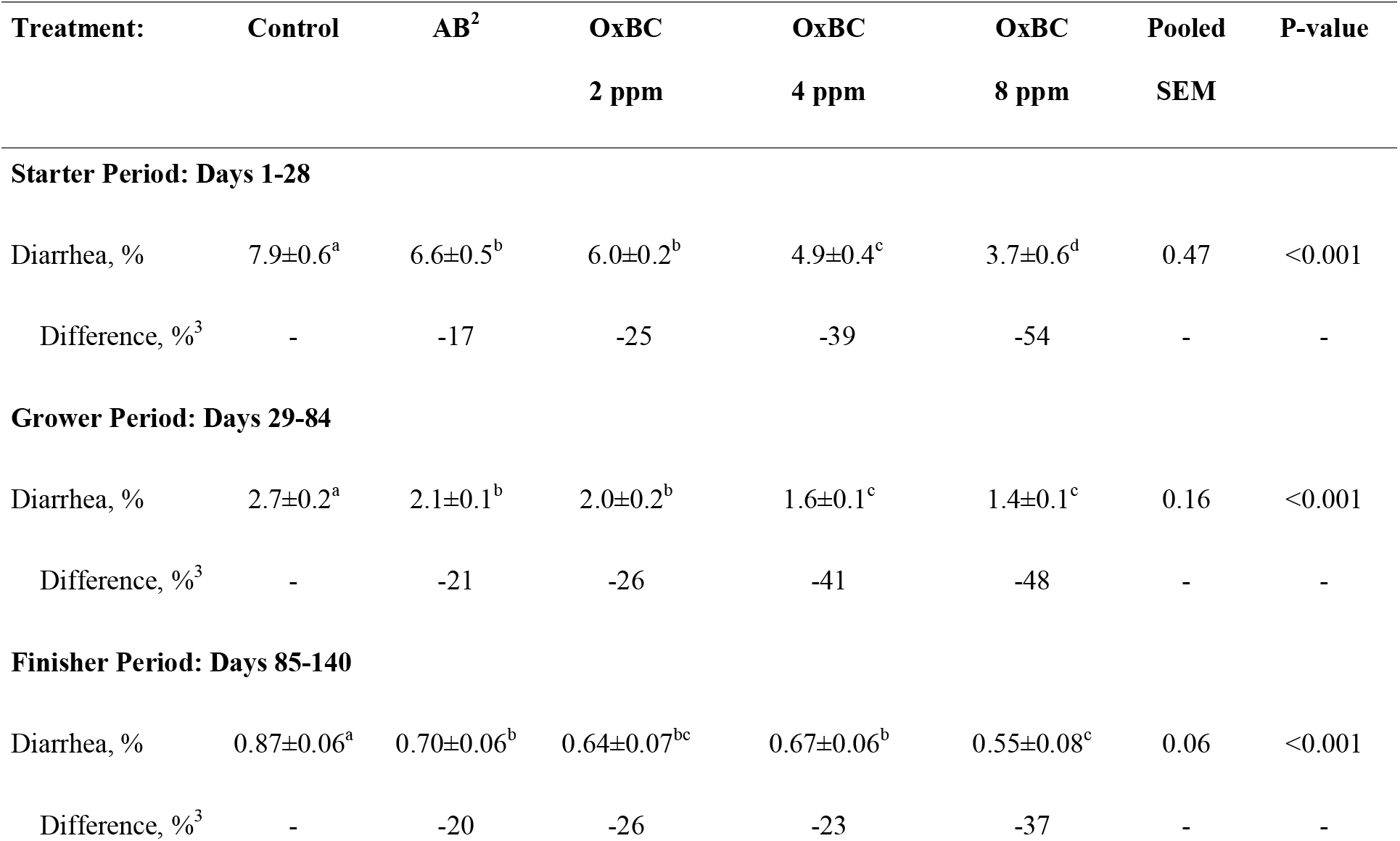
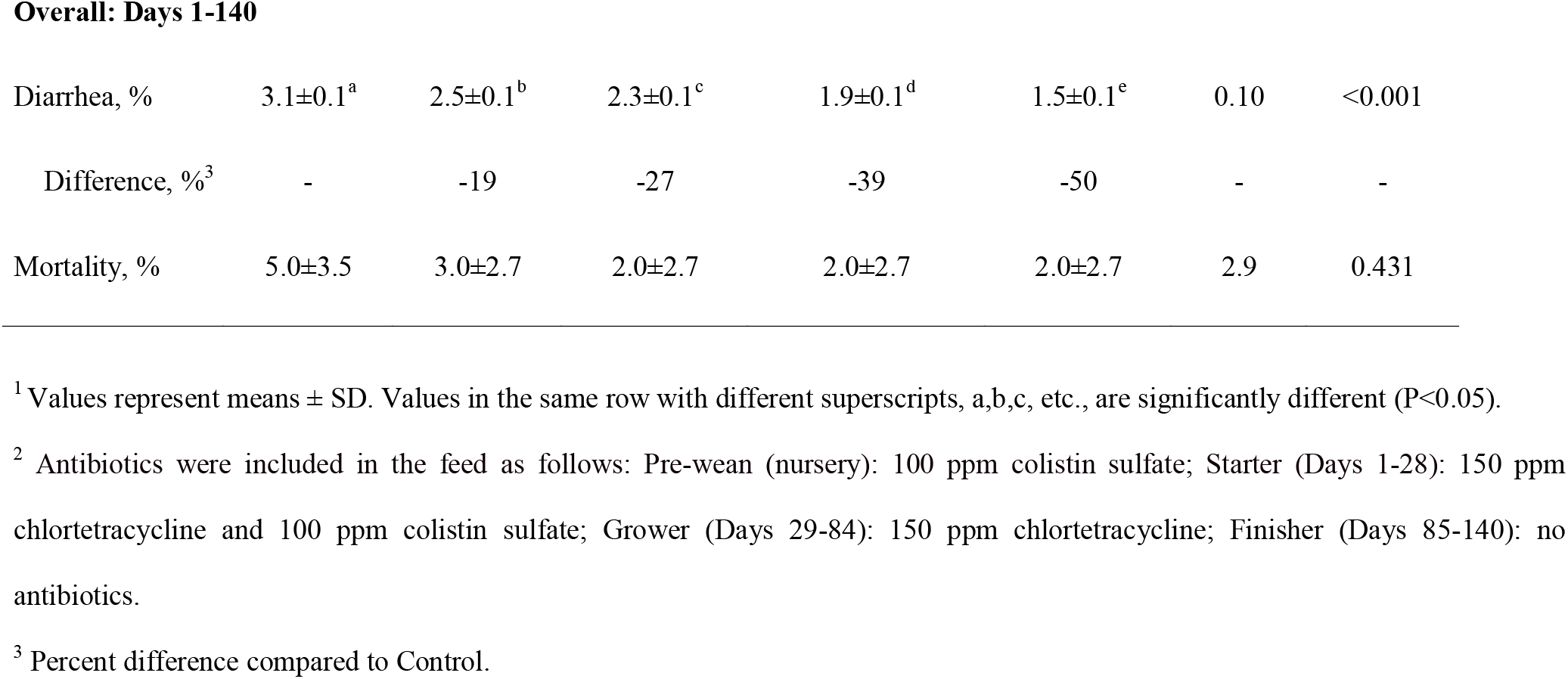
Effect of dietary OxBC on diarrhea incidence and overall mortality of pigs in the Starter, Grower, Finisher and Overall periods.^1^

| **Treatment:** | **Control** | **AB<sup>2</sup>** | **OxBC** | **OxBC** | **OxBC** | **Pooled** | **P-value** |
| --- | --- | --- | --- | --- | --- | --- | --- |
|  |  |  | **2 ppm** | **4 ppm** | **8 ppm** | **SEM** |  |
| **Starter Period: Days 1-28** |  |  |  |  |  |  |  |
| Diarrhea, % | 7.9±0.6<sup>a</sup> | 6.6±0.5<sup>b</sup> | 6.0±0.2<sup>b</sup> | 4.9±0.4<sup>c</sup> | 3.7±0.6<sup>d</sup> | 0.47 | <0.001 |
| Difference, %<sup>3</sup> | - | -17 | -25 | -39 | -54 | - | - |
| **Grower Period: Days 29-84** |  |  |  |  |  |  |  |
| Diarrhea, % | 2.7±0.2<sup>a</sup> | 2.1±0.1<sup>b</sup> | 2.0±0.2<sup>b</sup> | 1.6±0.1<sup>c</sup> | 1.4±0.1<sup>c</sup> | 0.16 | <0.001 |
| Difference, %<sup>3</sup> | - | -21 | -26 | -41 | -48 | - | - |
| **Finisher Period: Days 85-140** |  |  |  |  |  |  |  |
| Diarrhea, % | 0.87±0.06<sup>a</sup> | 0.70±0.06<sup>b</sup> | 0.64±0.07<sup>bc</sup> | 0.67±0.06<sup>b</sup> | 0.55±0.08<sup>c</sup> | 0.06 | <0.001 |
| Difference, %<sup>3</sup> | - | -20 | -26 | -23 | -37 | - | - |
| **Overall: Days 1-140** |  |  |  |  |  |  |  |
| Diarrhea, % | 3.1±0.1<sup>a</sup> | 2.5±0.1<sup>b</sup> | 2.3±0.1<sup>c</sup> | 1.9±0.1<sup>d</sup> | 1.5±0.1<sup>e</sup> | 0.10 | <0.001 |
| Difference, %<sup>3</sup> | - | -19 | -27 | -39 | -50 | - | - |
| Mortality, % | 5.0±3.5 | 3.0±2.7 | 2.0±2.7 | 2.0±2.7 | 2.0±2.7 | 2.9 | 0.431 |

<sup>1</sup> Values represent means ± SD. Values in the same row with different superscripts, a,b,c, etc., are significantly different (P<0.05).
<sup>2</sup> Antibiotics were included in the feed as follows: Pre-wean (nursery): 100 ppm colistin sulfate; Starter (Days 1-28): 150 ppm chlortetracycline and 100 ppm colistin sulfate; Grower (Days 29-84): 150 ppm chlortetracycline; Finisher (Days 85-140): no antibiotics.
<sup>3</sup> Percent difference compared to Control.

The occurrence of diarrhea during the Grower and Finisher periods was markedly less compared to the Starter period. Despite the lower background incidence of diarrhea in these later growth phases, OxBC still continued to provide significant and dose-dependent protection from the condition.

Overall, animals in the Control group had the highest incidence, whereas animals in the 8 ppm OxBC group had the lowest incidence. OxBC significantly and dose-dependently reduced the incidence relative to both the Control and AB groups (P<0.001).

Mortality was relatively low, overall, with few pigs dying during the study. The lowest number of deaths occurred in the OxBC groups, but differences were not significant between any of the groups.

### Trial 2: Pre-and post-wean piglet study

#### Stage 1 Pre-weaning (Days 1-21)

The effects of OxBC and antibiotics on the growth performance of pre-weaning piglets are provided in Table 5.

**Table 5.** BW, ADG, ADFI and F/G of pre-weaning piglets in Stage 1 of Trial 2 (Days 1-21)^1,2^

| Treatment: | Control | AB<sup>3</sup> | OxBC | OxBC | OxBC | OxBC | SEM | P-value |
| --- | --- | --- | --- | --- | --- | --- | --- | --- |
|  |  |  | 2 ppm | 4 ppm | 8 ppm | 16 ppm |  |  |
| BW Day 0 | 2.45±0.07 | 2.44±0.09 | 2.43±0.08 | 2.45±0.10 | 2.46±0.05 | 2.45±0.07 | 0.078 | 0.976 |
| Difference, %<sup>4</sup> | - | -0.4 | -0.8 | 0.0 | 0.8 | 0.4 |  |  |
| BW Day 21 | 6.88±0.07<sup>c</sup> | 7.21±0.07<sup>a</sup> | 6.92±0.17<sup>bc</sup> | 6.94±0.12<sup>bc</sup> | 6.98±0.08<sup>bc</sup> | 7.10±0.13<sup>ab</sup> | 0.11 | 0.001 |
| Difference, %<sup>4</sup> | - | 4.8 | 0.6 | 0.9 | 1.3 | 3.2 |  |  |
| ADG | 211±2.3<sup>c</sup> | 227±2.8<sup>a</sup> | 214±7.5<sup>bc</sup> | 214±2.9<sup>bc</sup> | 215±4.2<sup>bc</sup> | 222±7.8<sup>ab</sup> | 5.1 | 0.001 |
| Difference, %<sup>4</sup> | - | 8.1 | 1.9 | 1.9 | 2.4 | 5.2 |  |  |
| ADFI | 79.9±0.9 | 80.1±0.9 | 79.9±0.9 | 79.7±1.2 | 80.5±1.1 | 79.5±1.5 | 1.1 | 0.592 |
| Difference, %<sup>4</sup> | - | 0.9 | 0.1 | -0.3 | 0.9 | -0.4 |  |  |
| Feed/Gain | 0.38±0.01<sup>a</sup> | 0.35±0.01<sup>c</sup> | 0.37±0.01<sup>ab</sup> | 0.37±0.01<sup>ab</sup> | 0.37±0.01<sup>ab</sup> | 0.36±0.02<sup>bc</sup> | 0.01 | 0.001 |
| Difference, %<sup>4</sup> | - | -7.9 | -2.6 | -2.6 | -2.6 | -5.3 |  |  |
| Intake of Active<sup>5</sup> | 0 | 8.05 | 0.16 | 0.32 | 0.64 | 1.27 | - | - |

<sup>1</sup> Values represent means ± SD. Values in the same row with different superscripts, a,b,c, are significantly different (P<0.05).
<sup>2</sup> Units: BW: kg/head; ADG, ADFI: g/head/d; Intake of Active: mg/d.
<sup>3</sup> The AB diet contained 100 ppm colistin sulfate.
<sup>4</sup> Percent difference compared to Control.
<sup>5</sup> Average daily intake (mg) of antibiotic or OxBC.

Inclusion of OxBC in the creep feed resulted in a dose-dependent trend towards increased ADG and BW that reached significance for 16 ppm OxBC compared to the Control over the 21 days of Stage 1. The ADG and BW of the AB group were also higher than the Control. Antibiotics and 16 ppm OxBC improved the F/G ratio compared to the Control, while the ADFI was not significantly different across the 6 treatment groups (P>0.05).

The very low F/G ratio observed for all groups is attributed to the small amounts of feed ingested by the piglets, which were still mainly consuming milk from the sows.

No evidence of diarrhea or mortality was observed in Stage 1.

#### Trial 2: Stage 2 (Days 22-49)

The effects of OxBC and antibiotics on BW, ADG, ADFI and F/G for weaned piglets are shown in Table 6.

**Table 6.**
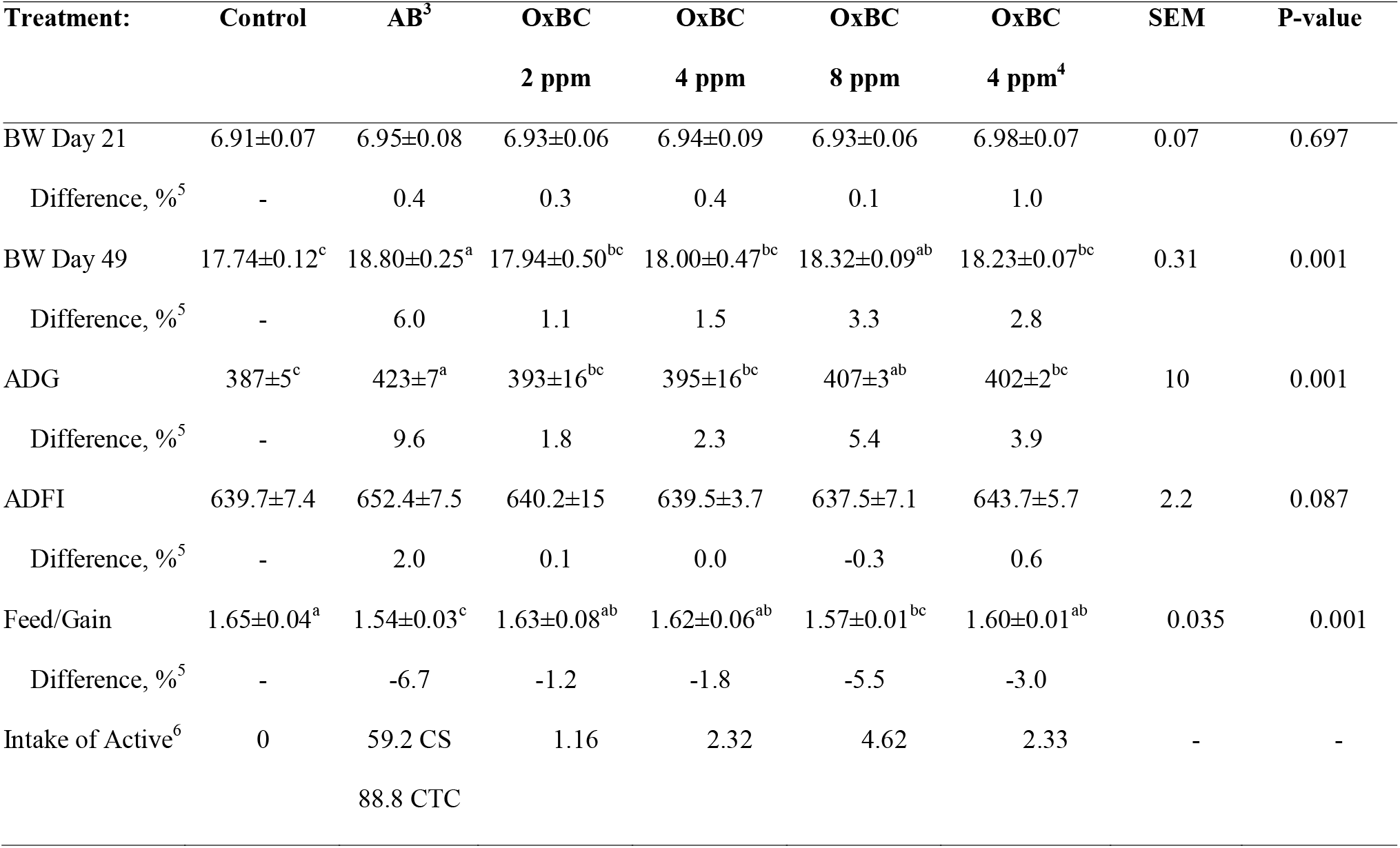
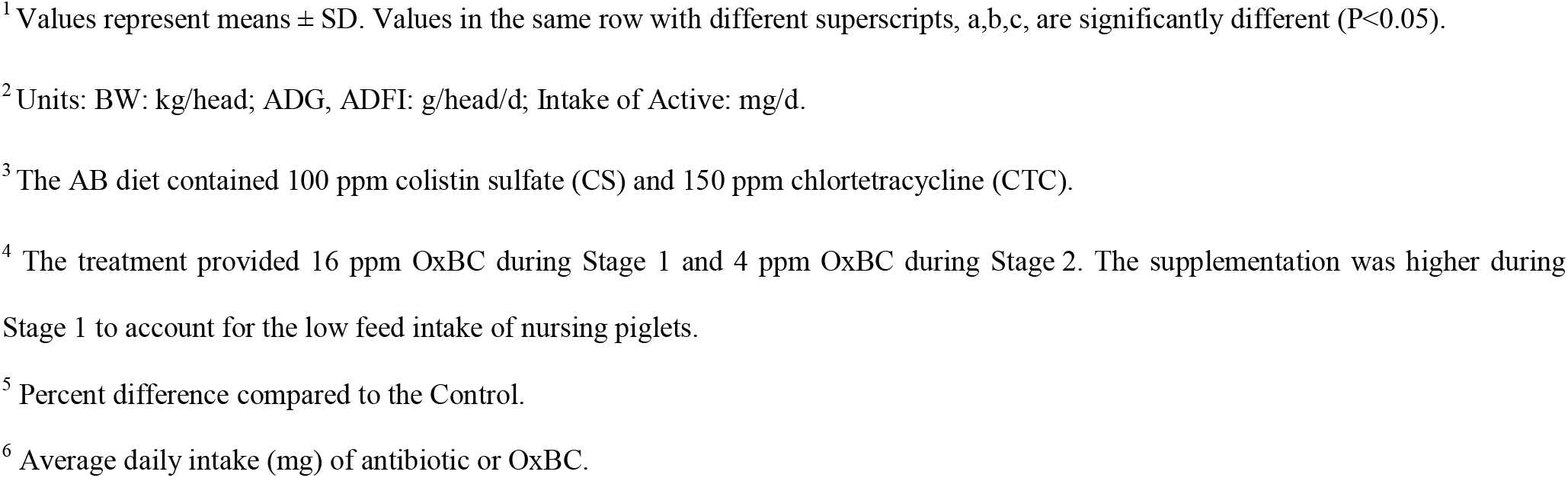
BW, ADG, ADFI and F/G of weaned piglets in Stage 2 of Trial 2 (Days 22-49).^1,2^

| **Treatment:** | **Control** | **AB<sup>3</sup>** | **OxBC** | **OxBC** | **OxBC** | **OxBC** | **SEM** | **P-value** |
| --- | --- | --- | --- | --- | --- | --- | --- | --- |
|  |  |  | **2 ppm** | **4 ppm** | **8 ppm** | **4 ppm<sup>4</sup>** |  |  |
| BW Day 21 | 6.91±0.07 | 6.95±0.08 | 6.93±0.06 | 6.94±0.09 | 6.93±0.06 | 6.98±0.07 | 0.07 | 0.697 |
| Difference, %<sup>5</sup> | - | 0.4 | 0.3 | 0.4 | 0.1 | 1.0 |  |  |
| BW Day 49 | 17.74±0.12<sup>c</sup> | 18.80±0.25<sup>a</sup> | 17.94±0.50<sup>bc</sup> | 18.00±0.47<sup>bc</sup> | 18.32±0.09<sup>ab</sup> | 18.23±0.07<sup>bc</sup> | 0.31 | 0.001 |
| Difference, %<sup>5</sup> | - | 6.0 | 1.1 | 1.5 | 3.3 | 2.8 |  |  |
| ADG | 387±5<sup>c</sup> | 423±7<sup>a</sup> | 393±16<sup>bc</sup> | 395±16<sup>bc</sup> | 407±3<sup>ab</sup> | 402±2<sup>bc</sup> | 10 | 0.001 |
| Difference, %<sup>5</sup> | - | 9.6 | 1.8 | 2.3 | 5.4 | 3.9 |  |  |
| ADFI | 639.7±7.4 | 652.4±7.5 | 640.2±15 | 639.5±3.7 | 637.5±7.1 | 643.7±5.7 | 2.2 | 0.087 |
| Difference, %<sup>5</sup> | - | 2.0 | 0.1 | 0.0 | -0.3 | 0.6 |  |  |
| Feed/Gain | 1.65±0.04<sup>a</sup> | 1.54±0.03<sup>c</sup> | 1.63±0.08<sup>ab</sup> | 1.62±0.06<sup>ab</sup> | 1.57±0.01<sup>bc</sup> | 1.60±0.01<sup>ab</sup> | 0.035 | 0.001 |
| Difference, %<sup>5</sup> | - | -6.7 | -1.2 | -1.8 | -5.5 | -3.0 |  |  |
| Intake of Active<sup>6</sup> | 0 | 59.2 CS | 1.16 | 2.32 | 4.62 | 2.33 | - | - |
|  |  | 88.8 CTC |  |  |  |  |  |  |

<sup>1</sup> Values represent means ± SD. Values in the same row with different superscripts, a,b,c, are significantly different (P<0.05).
<sup>2</sup> Units: BW: kg/head; ADG, ADFI: g/head/d; Intake of Active: mg/d.
<sup>3</sup> The AB diet contained 100 ppm colistin sulfate (CS) and 150 ppm chlortetracycline (CTC).
<sup>4</sup> The treatment provided 16 ppm OxBC during Stage 1 and 4 ppm OxBC during Stage 2. The supplementation was higher during Stage 1 to account for the low feed intake of nursing piglets.
<sup>5</sup> Percent difference compared to the Control.
<sup>6</sup> Average daily intake (mg) of antibiotic or OxBC.

At the beginning of Stage 2, a subset of select piglets from each treatment group was randomized within treatment groups to continue the trial. Piglets were selected such that the average BW among the six treatment groups did not differ on Day 22 (the beginning of Stage 2). After 28 days (Day 49, the end of Stage 2), average BWs were highest for the AB group followed by the four OxBC groups and the Control. Increases in average BW reached statistical significance for both the AB and the 8 ppm OxBC groups (P<0.001).

The increases in BW were reflected in the ADG. The AB group had the highest ADG, which was significantly higher than the Control (P<0.001). ADG showed a trend towards a dose-dependent increase in the OxBC groups, reaching statistical significance for the 8 ppm OxBC group compared to the Control. However, there was no statistical difference in ADG across the four OxBC groups.

There were no significant differences in ADFI among the treatment groups. F/G ratios were significantly improved for both the AB group and the 8 ppm OxBC group compared to all other groups (P<0.001).

The effects on the incidence of diarrhea and mortality are given in Table 7. Diarrhea and mortality incidence were generally low in all groups, with the highest incidence of diarrhea and mortality being observed in the Control group. Treatment with 8 ppm OxBC or antibiotics decreased diarrhea incidence by 24% and 28%, respectively.

**Table 7.** Diarrhea incidence and mortality in weaned piglets during Stage 2 (Days 22-49)

| **Treatment:** | **Control** | **AB<sup>1</sup>** | **OxBC** | **OxBC** | **OxBC** | **OxBC** |
| --- | --- | --- | --- | --- | --- | --- |
|  |  |  | **2 ppm** | **4 ppm** | **8 ppm** | **4 ppm<sup>2</sup>** |
| Diarrhea, % | 4.7±0.8 | 3.4±1.1 | 4.6±1.3 | 4.1±0.9 | 3.6±1.1 | 3.9±0.8 |
| Difference,%<sup>3</sup> | - | -28 | -1 | -12 | -24 | -18 |
| Mortality, % | 5.0 | 0 | 3.4 | 3.4 | 0 | 0 |
| vs. Control, % | (100) | 0 | 67 | 67 | 0 | 0 |

<sup>1</sup> The AB diet contained 100 ppm colistin sulfate and 150 ppm chlortetracycline.
<sup>2</sup> The treatment provided 16 ppm OxBC during Stage 1 and 4 ppm OxBC during Stage 2. The supplementation was higher during Stage 1 to account for the low feed intake of nursing piglets.
<sup>3</sup> Percent difference compared to the Control.

No mortalities were recorded in the AB group, the 8 ppm OxBC group or the group fed 16 ppm OxBC in Stage 1 and 4 ppm OxBC in Stage 2. The differences in mortalities between the groups were not significant.

## Discussion

Results from the two trials reported here demonstrate that dietary supplementation with low parts-per-million levels of OxBC improves the health and productivity of pigs reared under commercial production conditions. In Trial 1, conducted over the entire 140-day post-wean grow-out period, inclusion of OxBC in the feed led to improvements in growth performance during each of the three stages of the production cycle and also significantly reduced the incidence of diarrhea in post-wean starter piglets. Results from the second trial confirm the benefits of dietary OxBC on growth performance of post-wean starter piglets and indicate that further benefits on piglet performance are possible when OxBC is added to the transition (creep) feed of nursing piglets prior to weaning.

In Trial 1, the observation that the benefits of OxBC were most apparent during the starter period compared to the grower and finisher periods is not surprising, given that OxBC plays a role in supporting immunity (Burton et al., 2014; Duquette et al., 2014; Johnston et al., 2014) and the fact that young piglets experience high levels of stress associated with weaning and that they are immunologically immature. The highest incidence rates of diarrhea occurred during the starter period, consistent with the well-known susceptibility of young piglets to post-wean diarrhea (Rhouma et al., 2017). The reduced incidence of diarrhea observed in the OxBC supplemented groups relative to controls is consistent with the immune priming actions of OxBC, which center upon an ability to increase the level of pathogen pattern recognition receptors (PPRR) and enhanced down-stream innate immune response to receptor activation (Johnston et al., 2014).

More specifically, OxBC has been shown to increase the level of toll-like receptor subtypes 2 and 4 (TLR-2, TLR-4) and their coreceptor CD14 in both *in vitro* and *in vivo* studies (Johnston et al., 2014). TLR-4 and CD14 are well-characterized co-receptors responsible for recognizing the lipopolysaccharide (LPS) moiety present in gram negative bacteria, such as *E. coli*, while TLR-2 recognizes the lipoteichoic acid (LTA) moiety present in gram positive bacteria, such as *Clostridium perfringens* (Dessing et al., 2008; Park and Lee, 2013; Takehara, 2018). Binding of bacterial LPS or LTA to the TLR-4/CD14 or TLR-2 receptor triggers an innate immune response aimed at clearing the bacteria (Mukherjee et al., 2016; Vijay, 2018).

A study comparing *E. coli* resistant versus susceptible lineages of pigs found that the genetic basis for the resistance to *E. coli* colonization of the gut was higher levels of CD14 expression (Wu et al., 2016). The authors concluded that the TLR-4/CD14 pathway plays a significant role in reducing *E. coli* colonization and the incidence of diarrhea in piglets. While we did not measure intestinal receptor or *E. coli* levels in the current trials, an increase in intestinal CD14 and TLR-4 content (as we have observed in mice fed OxBC (Johnston et al., 2014)) and the subsequent reduction in *E. coli* colonization offers a plausible explanation for the reduced incidence of diarrhea observed in the OxBC-supplemented piglets. Results from an infectious challenge study in broilers, where dietary supplementation with OxBC reduced *Clostridium perfringens* levels by 2 to 3 log units, are consistent with the concept that OxBC supports the intestinal immune system and allows the host to resist pathogen colonization (Kang et al., 2018).

The improvement in clinical health (reduced incidence of diarrhea) likely explains, in part, the improved growth performance of piglets in the OxBC-supplemented versus control groups during the Starter period in Trial 1. As would be expected for older, more immunocompetent pigs, there were fewer incidences of diarrhea in grower and finisher pigs compared to starter pigs across all treatment groups in the first trial. While there tended to be a lower incidence of diarrhea in the OxBC supplemented groups compared to the control during the grower and finisher periods, it seems unlikely that these small reductions in disease incidence would explain the improved growth performance observed in the OxBC groups during the latter stages of the trial. Instead, the ability of OxBC to reduce inflammatory tone represents the most plausible mechanism to explain the improved growth of older pigs in the OxBC groups.

The intestine has been described by some authors as a tissue in a constant state of controlled inflammation, and there is a need for organisms to tightly control the level of intestinal inflammation (Biancone et al., 2002; Niewold, 2007). Commercial production practices are known to further exacerbate the inflammatory state of the intestine, as stresses associated with weaning, *ad libitum* feeding of high energy diets, and the presence of anti-nutritional components in the diet all contribute to increased inflammation of the gut (Pluske et al., 2018). The contribution that reduced inflammation plays in improving growth performance is further highlighted by the fact that an anti-inflammatory action has been proposed as an underlying mechanism contributing to the growth promoting effects of subtherapeutic antibiotic growth promoters (Niewold, 2007).

Results from studies in an experimental model of bovine respiratory disease (BRD) and in gestating/lactating sows demonstrate an anti-inflammatory action for OxBC. In the experimental BRD trial, animals that received dietary supplementation with OxBC had a significant increase in anti-inflammatory or pro-resolution activity in the lung (Duquette et al., 2014) while OxBC-supplementation in gestating/lactating sows led to reductions in the concentration of pro-inflammatory cytokines, TNFα and IL-8 in colostrum and milk (Chen et al., 2020). Reduced inflammation may also be partly responsible for the observed reduction in intestinal lesion severity in *C. perfringens-challenged* broilers (Kang et al., 2018).

The performance benefits of OxBC in weaned piglets observed in Trial 1 are supported by the results from Trial 2 in which inclusion of OxBC in the Starter feed led to an apparent dose-dependent trend for improved growth performance and feed efficiency, with the highest inclusion level of OxBC (8 ppm) reaching significance. In terms of clinical health, the results from Trial 2 indicate that, while the incidences of diarrhea were numerically lower in the antibiotic and OxBC supplemented groups compared to the control, there was no significant treatment effect (Table 7). There were considerably fewer cases of diarrhea observed in the control group in Trial 2 (4.7%) compared to Trial 1 (7.9%). This difference in the background incidence of diarrhea in the two trials likely reflects a lower level of endemic pathogen pressure during Trial 2 relative to Trial 1. With a relatively low incidence of diarrhea observed in the control group in Trial 2, further treatment-induced reductions may not have been possible. Thus, in Trial 2 the benefits of OxBC on growth performance of post-wean piglets is more plausibly due to anti-inflammatory actions rather than a reduction in the incidence of diarrhea. It is interesting to note that the magnitude of the performance benefits in piglets was much larger in Trial 1 compared to Trial 2. The difference in the magnitude of the benefits may have been due to a combined benefit of reducing diarrhea and background inflammation in the gut in Trial 1, while in the second trial, where there was lower endemic pathogen pressure, only the anti-inflammatory activity may have factored in the benefit.

The results from the pre-wean period (Stage 1) of Trial 2 indicate that inclusion of OxBC in transition or creep feed can also improve weight gain and body weight of nursing piglets. OxBC appeared to dose-dependently increase ADG and body weight at weaning with the highest inclusion rate of 16 ppm OxBC reaching statistical significance (Table 6). The lack of a statistically detectable effect at the lower OxBC inclusion rates likely reflects the very low amounts of OxBC received by nursing piglets that consumed relatively little feed. Given the apparent dose-dependent improvements in growth performance, it would be useful to investigate higher inclusion rates of OxBC in transition feed or OxBC-supplementation of milk replacer. There were no mortalities or incidences of diarrhea observed during the pre-wean stage of Trial 2, consistent with the presence of a relatively low level of endemic pathogen pressure.

Overall, the results of the two trials reported here demonstrate the benefits of dietary supplementation with OxBC on growth performance, feed efficiency, and the health of pre- and post-weaned piglets. These findings have significance for both the commercial feed industry and the field of nutrition. For the feed industry, the findings support the application of OxBC as an effective alternative to antibiotic growth promoters. The performance and health benefits obtained with OxBC were comparable to those of the antibiotic growth promoters in both trials. For the field of nutrition, the results lend further support to the identification of the copolymers present in OxBC as a source of the benefits of dietary β-carotene. The concept that β-carotene is a source of beneficial compounds beyond vitamin A is not new; however, there is a dearth of reproducible evidence as to the identity and biological actions of these compounds. A scientific panel of the European Food Safety Association (EFSA) recently concluded there was insufficient scientific evidence to support a proposed benefit of β-carotene, as a feed additive, on host immunity (EFSA, 2012). The EFSA panel cited an inconsistency of results with β-carotene as the major problem. The inconsistent results cited by EFSA likely stem from the fact that the proposed immunological benefits arise not from vitamin A or a direct action of β-carotene itself but rather from the presence of copolymers, which were unknown at the time. Dietary supplementation with β-carotene potentially provides variable levels of copolymers, depending upon the extent of adventitious oxidation. β-Carotene’s susceptibility to oxidation is recognized by manufacturers who nowadays formulate the product to help prevent its oxidation.

In the trials reported here, the source of the β-carotene copolymers is a commercial product manufactured by a highly reproducible process under stringent quality control standards. However, the copolymers have also been shown to exist naturally at varying levels in various food and feedstuffs (Burton et al., 2016; Schaub et al., 2017). The presence of naturally occurring copolymers may provide an explanation, at least in part, for the health and productivity benefits associated with the use of certain traditional forages. For example, alfalfa, a forage rich in β-carotene, has been proposed to contain unidentified growth factors that benefit the growth and reproductive performance of various species (Scott et al., 1953; Lakhanpal et al., 1966; Hertrampf and Piedad-Pascual, 2000). The mechanisms of action of the so-called unidentified growth factors remains largely unknown; however the presence of naturally occurring copolymers arising from oxidation of β-carotene when alfalfa is dried (Burton et al., 2016) represents one possible growth factor.

We propose that β-carotene is the source of two classes of compounds that are biologically active and which both benefit health, the first being vitamin A and the second being β-carotene oxygen copolymers. Driven by the desire to reduce costs, the modern feed industry has largely replaced its use of β-carotene, whether synthetic or natural, with synthetic vitamin A. Removal of β-carotene, particularly natural sources, such as alfalfa, from diets has resulted in the simultaneous, albeit unintended, removal of the naturally present β-carotene copolymers and their associated benefits. The positive effects on growth performance and health observed with OxBC in these trials coupled with similar findings in previous trials in swine (Hurnik et al., 2011; Chen et al., 2020), poultry (Kang et al., 2018) and dairy calves (Duquette et al., 2014) demonstrate that re-introduction of very low amounts of β-carotene oxygen copolymer into feeds can beneficially support the health and productivity of livestock and substantially reduce the need for in-feed antibiotics.

## Abbreviations

OxBC: oxidized β-carotene
AB: antibiotic diet
ADG: average daily growth
ADFI: average daily feed intake
BW: body weight
CD14: cluster of differentiation receptor 14
CP: crude protein
DCP: dicalcium phosphate
DDGS: dried distillers grains with solubles
F/G: feed/gain
ME: metabolizable energy
NE: net energy
OxBC: fully oxidized β-carotene
PPRR: pathogen pattern recognition receptor
PRRS: porcine reproductive and respiratory syndrome
TLR: toll-like receptor

